# A new reporter cell line for studies with proteasome inhibitors in *Trypanosoma brucei*

**DOI:** 10.1101/364646

**Authors:** Danielle MN Moura, Osvaldo P de Melo Neto, Mark Carrington

## Abstract

A *Trypanosoma brucei* cell line is described that produces a visual readout of proteasome activity. The cell line contains an integrated transgene encoding an ubiquitin-green fluorescent protein (GFP) fusion polypeptide responsive to the addition of proteasome inhibitors. A modified version of *T. brucei* ubiquitin unable to be recognized by deubiquitinases (UbG76V) was fused to eGFP and constitutively expressed. The fusion protein is unstable but addition of the proteasome inhibitor lactacystin stabilizes it and leads to visually detectable GFP. This cell line can be widely used to monitor the efficiency of inhibitor treatment through detection of GFP accumulation in studies involving proteasome-mediated proteolysis, screening of proteasome inhibitors or other events related to the ubiquitin-proteasome pathway.

## Main text

The proteasome is a multi-catalytic ATP-dependent protease complex that plays a central role in the ubiquitin-mediated proteolysis, the major pathway for regulated degradation of multiple protein targets including cytosolic, nuclear and membrane polypeptides in all eukaryotic organisms^1^. The process of ubiquitination is mediated by three enzymes (E1, E2 and E3) that act in series to generate an isopeptide bond between the carboxyl group of the C-terminal glycine of ubiquitin and the amino group on the side chain of a lysine residue on the substrate. This can then result in degradation of the targeted protein by the proteasome whereas the ubiquitin is recycled following the action of deubiquitinases^2^.

The ubiquitin-proteasome system has emerged as a therapeutic target for diverse pathologies such as cancer, neurodegenerative diseases, immune diseases and infections, including those caused by parasites^3^. Proteasome inhibitors are structurally diverse and can interact directly or allosterically with the proteasome active site(s), and can be reversible or irreversible^4^. Lactacystin, a β-lactone precursor from natural source, is an example of a potent and specific inhibitor of the proteasome proteolytic activity that binds irreversibly to the catalytic threonines found in the active sites of the proteasome β-subunits^5,6^.

*Trypanosoma brucei* and *T. cruzi* are the causative agents of African Trypanosomiasis and Chagas disease, respectively, widespread tropical diseases that can be fatal if not treated. Lactacystin as well as other compounds, such as MG132, have been shown to inhibit proteasome activity in both *T. brucei* and *T. cruzi*, and studies using these compounds have helped to clarify the role of proteasomes in cell proliferation and differentiation in these pathogens^7–9^. Recently, studies have tested inhibitors of the kinetoplastid proteasome, including the molecule GNF6702, which showed selective effect *in vivo*, corroborating the idea that the proteasome is a potential target for treatment of infections caused by parasites^10,11^.

Inhibitory concentrations of lactacystin has been determined in *Trypanosoma* species^8,12^, however the time taken for lactacystin to cause primary effects has not been investigated in most studies in trypanosomatids and long incubations, usually over 10 hours, are used based on protocols developed for mammalian cells. These long incubations can make it difficult to distinguish primary and secondary effects of proteasome inhibition. To circumvent the problems above and to monitor the *T. brucei* response to lactacystin, we have produced a *T. brucei* reporter cell line based on the fusion of ubiquitin to a reporter fluorescent protein, an approach first developed in HeLa cells^13^. The *T. brucei* ubiquitin gene (Tb927.11.9920) encodes a polyubiquitin with nine tandem ubiquitin repeats. DNA fragments encoding single ubiquitin polypeptides (76 amino acids) were amplified using PCR, one containing an open reading frame (ORF) encoding wild type ubiquitin and a second designed to produce a mutation in the C-terminal amino acid of ubiquitin sequence, changing it from a glycine to a valine (G76V). Both PCR reactions also added 39 nucleotides encoding a 13 amino acid extension to the C-terminus of the wild type or G76V ubiquitin. The purpose of this extension was to meet the requirements for ubiquitin recognition and cleavage by deubiquitinases (Figure 1A). Each PCR product was cloned between the *EcoR*I and *Hind*III sites of a modified version of p3605^14^, which contains an eGFP ORF in a construct designed to insert into the tubulin locus by homologous recombination (Figure 1B). The result was two plasmids containing a transgene encoding a chimeric protein, either Ub-GFP (plasmid p4596) or Ub(G76V)-GFP (plasmid p4595). The mutation in Ub(G76V)-GFP means it is not cleaved by deubiquitinases and instead is degraded by the proteasome. Ub-GFP represented a control in which the polypeptide is cleaved by a deubiquitinase releasing stable GFP. The final constructs, p4595 and p4596, were cut with the restriction enzyme PacI and transfected into a procyclic form *Trypanosoma brucei* Lister427 pSMOx cell line^15^. Selection with 15 μg/ml geneticin (G418) in SDM-79 culture medium was used to select the respective cell lines, Lister427 pSMOx p4595 and and Lister427 pSMOx p4596.

**Figure 1.**
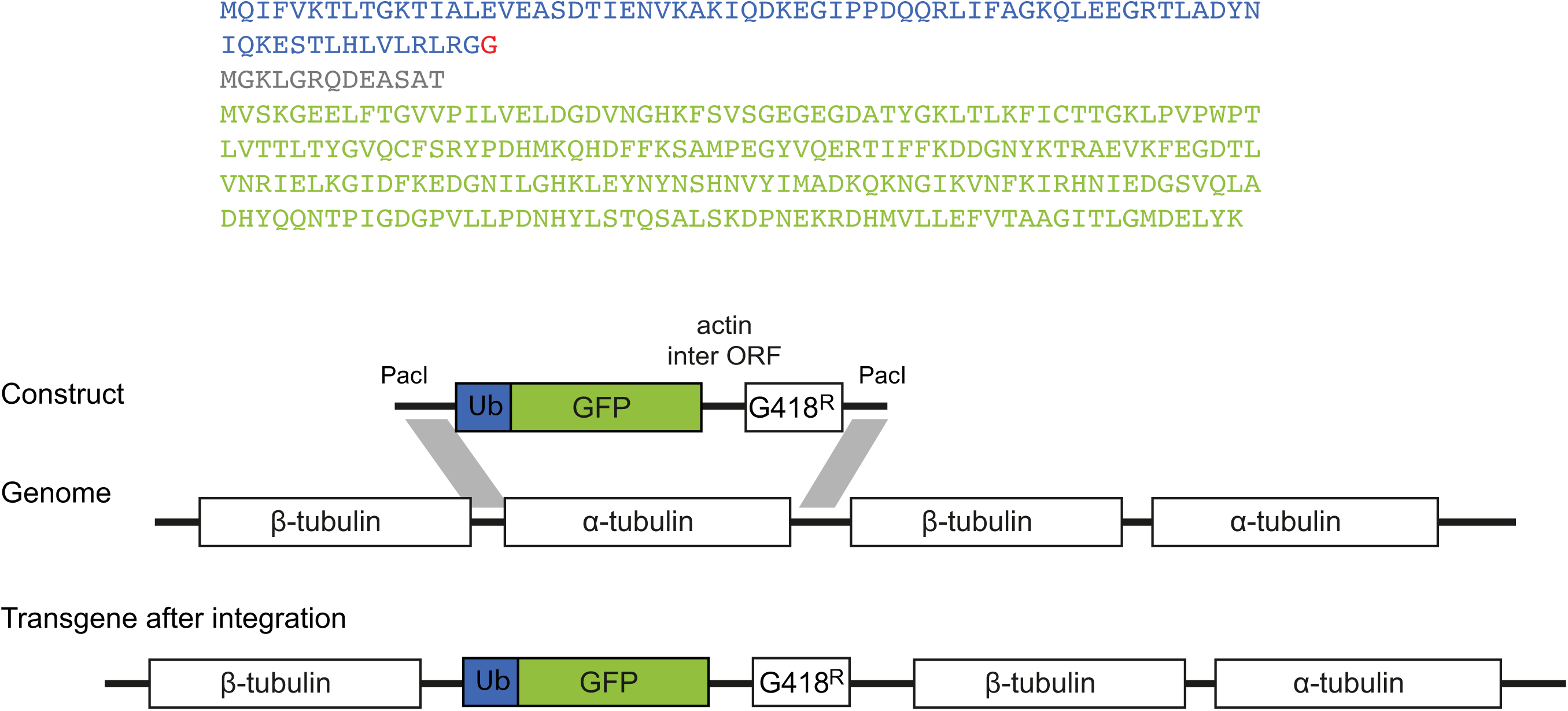
Establishing the reporter cell lines. A) Sequence of the ubiquitin-green fluorescent proteins encoded by the transgenes. Ubiquitin is shown in blue with glycine 76, mutated to valine in the G76V variant, indicated in red; the linker is coloured in grey and the green fluorescent protein coding sequence in green. B) Representation of the insertion of the transgene construct into the tubulin locus by targeted homologous recombination. The construct resulted in the expression of a transgene mRNA with an alpha tubulin 5’UTR, transgene ORF and actin 3’UTR. Transcription was a result of read through by RNA polymerase II.

The cell lines were then incubated with 5 μM lactacystin in culture medium during log phase growth and GFP levels measured over a time course by western blotting, fluorescence microscopy and flow cytometry. The cell line containing the Ub(G76V)-GFP transgene had little GFP fluorescence before lactacystin addition consistent with being targeted for proteasomal degradation. After lactacystin addition, GFP fluorescence became apparent (Figure 2A and B) and GFP with a molecular weight of ~35 kDa was detected by western blotting, consistent with the Ub(G76V)-GFP fusion protein (Figure 2C). In contrast, the cell line containing the Ub-GFP transgene constitutively expressed GFP, detected by fluorescence microscopy and as a 25 kDa polypeptide by western blot (Figure 2A, B and C). The action of lactacystin in accumulating Ub(G76V)-GFP occurred in the first hours of incubation (up to 8h) (Figure 2A and B), with GFP being detected by fluorescence microscopy as early as 2 hours of lactacystin treatment (data not shown).

**Figure 2.**
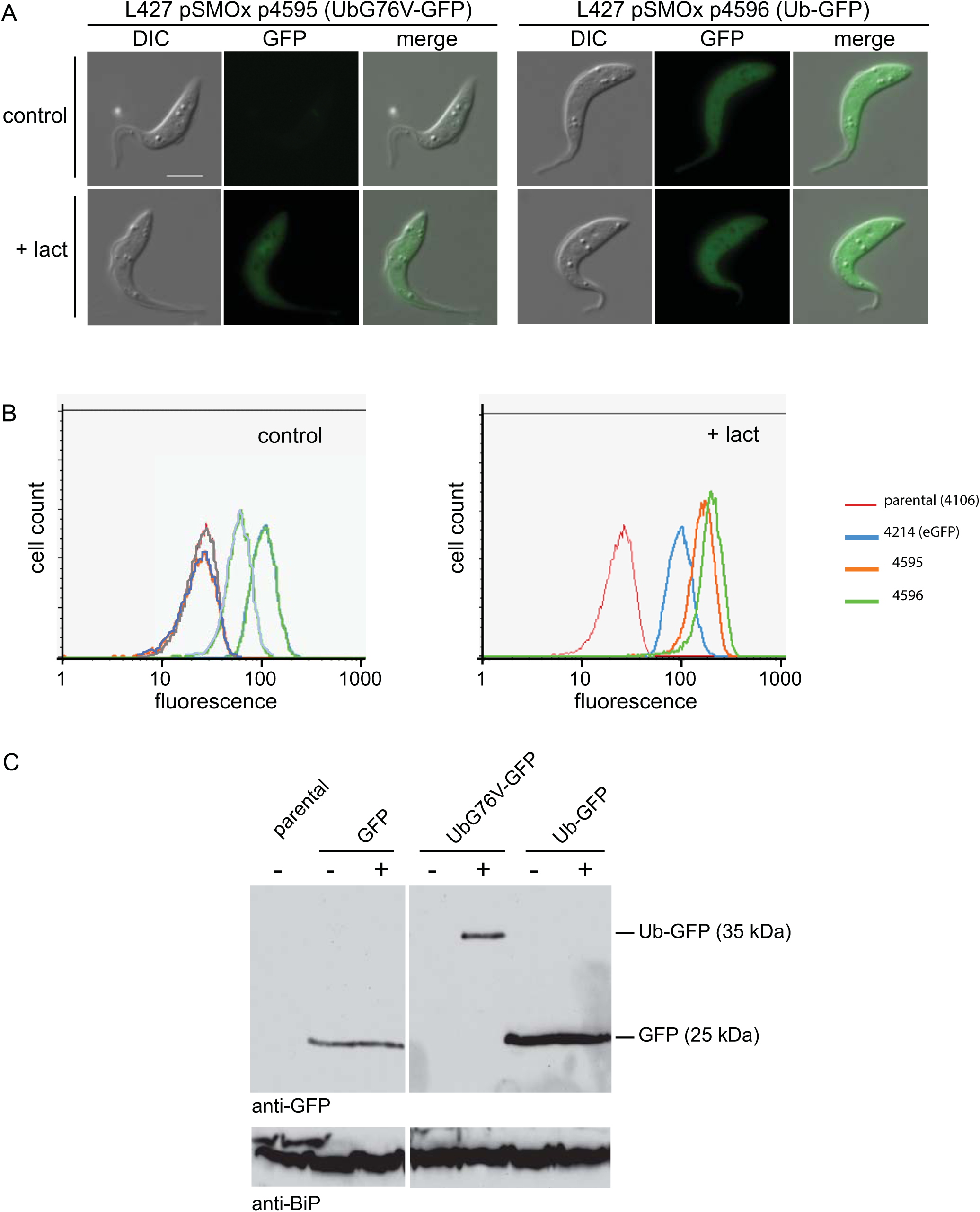
Detection of GFP expression after lactacystin treatment. A) Fluorescence microscopy detection of GFP expression before and after incubation with 5 μM lactacystin for 6 hours. White scale bar = 5 μm. B) Flow cytometry analysis of GFP expression before and after addition of 5 μM lactacystin for 4 hours. For the flow cytometry experiments, a parental cell line Lister427 pSMOx and Lister 427 p4214, a cell line expressing an untagged eGFP from the tubulin locus, were used as negative and positive control, respectively. C) GFP expression in different cell lines detected by western blotting using anti-GFP before or after incubation for 8 hours with 5 μM lactacystin as indicated. Cell lysate equivalent to 2×10^6^ cells were loaded in each lane and detection of the chaperone BiP was used as loading control.

The readily detection of accumulated GFP in cells expressing Ub(G76V)-GFP by either fluorescence microscopy or flow cytometry after treatment with lactacystin indicates that the cell line is an excellent indicator for proteasome inhibition. It is a convenient tool for further studies involving the ubiquitin-proteasome pathway and screening of new proteasomal inhibitors. The plasmids are available from the authors upon request.

## Acknowledgements

This work was funded by the Brazilian funding agencies FACEPE (ATP-0018-2.02/13) and Capes (PNPD 23038.007656/2011-92) and the Wellcome Trust (085956/Z/08/Z).

## Abbreviations

eGFP: enhanced green fluorescent protein
ub: ubiquitin
lact: lactacystin
ORF: open reading frame

